## Supplementary Information for "IDENTIFYING PLANT GENES SHAPING MICROBIOTA COMPOSITION IN THE BARLEY RHIZOSPHERE"

[1]University of Dundee, Plant Sciences, School of Life Sciences, Dundee, UK; [2]University of Dundee, Computational Biology, School of Life Sciences, Dundee, UK; [3]Mohammed VI Polytechnic University, Agrobiosciences Program, Plant & Soil Microbiome Subprogram, Bengurir, Morocco; [4]The James Hutton Institute, Invergowrie, UK; [5]Department of Biosciences and Territory, University of Molise, Campobasso, Italy; [6]Faculty of Science and Technology, Free University of Bozen-Bolzano, Bolzano, Italy; [7]Competence Centre for Plant Health, Free University of Bozen-Bolzano, Bolzano, Italy; [8]Scotland's Rural College, Edinburgh, UK; [9]Department of biological and environmental sciences and technologies, University of Salento, Lecce, Italy; [10]Institute of Agricultural and Food Science and Plant Breeding, Martin Luther University, Halle-Wittenberg, Germany; [11]These authors contributed equally: Carmen Escudero-Martinez and Max Coulter.

**Supplementary Table 1:** Genome-wide LOD threshold score calculated per individual taxa (phenotype) at ASV, genus and family level (1000 permutations, alpha=0.2). Source data are provided as a Source Data file.

| **ASV (alpha=0.2)** | | | | | | | | |
|---|---|---|---|---|---|---|---|---|
| ***Sphingomonas env.OPS 17*** | ***Allo.Neo.Para. Rhizobium*** | ***Pedobacter1*** | ***Paenibacillus1*** | ***Niastella*** | ***Paenibacillus2*** | ***Stenotrophomonas*** | ***Paenibacillus3*** | ***Streptomyces2*** |
| 6.92 | 8.92 | 8.8 | 3.19 | 6.18 | 3.31 | 15.0 | 3.68 | 3.93 |
| ***Lysobacter1*** | ***Asticcacaulis*** | ***Variovorax*** | ***Paenibacillus4*** | ***Bdellovibrio*** | ***Duganella*** | ***Holophaga*** | ***Vicinamibacterales*** | ***Sphingobacteri*** |
| 3.80 | 10.0 | 6.01 | 5.73 | 8.74 | 7.07 | 12.2 | 3.43 | 3.63 |
| ***Lysobacter2*** | ***Sorangium*** | ***Sphingopyxis*** | ***Chryseolinea*** | ***Tahibacter*** | ***Pedobacter2*** | ***Comamonadaceae*** | ***Chryseobacterium*** | ***Streptomyces3*** |
| 5.20 | 13.9 | 7.75 | 9.61 | 9.79 | 6.32 | 7.26 | 4.96 | 5.47 |
| ***Pedobacter3*** | ***Asticcacaulis1*** | ***Pseudomonas*** | ***Pseudomonas1*** | ***Streptomyces1*** | ***Lysobacter3*** | ***Sphingomonas*** | ***Nocardia*** | |
| 11.4 | 7.81 | 5.70 | 6.68 | 5.19 | 11.1 | 5.36 | 3.78 | |
| **Genus (alpha=0.2)** | | | | | | | | |
| ***Pedobacter*** | ***Paenibacillus*** | ***Ramlibacter*** | ***Rhodanobacter*** | ***Asticcacaulis*** | ***Thermomonas*** | ***Variovorax*** | ***Dokdonella*** | ***Chryseobacterium*** |
| 6.61 | 3.09 | 5.20 | 15.5 | 9.83 | 8.79 | 10.5 | 6.65 | 4.34 |
| ***Bdellovibrio*** | ***Microbacterium*** | ***P3OB.42*** | ***Holophaga*** | ***Sorangium*** | ***Chryseolinea*** | ***Tahibacter*** | ***Comamonadaceae*** | ***Streptomyces*** |
| 4.38 | 4.29 | 12.2 | 11.5 | 12.6 | 7.26 | 8.52 | 4.06 | 3.58 |
| **Family (alpha=0.2)** | | | | | | | | |
| ***Sphingobacteriaceae*** | ***Paenibacillaceae*** | ***Microscillaceae*** | ***Opitutaceae*** | ***Caulobacteraceae*** | ***Comamonadaceae*** | ***Chitinophagaceae*** | ***Alcaligenaceae*** | **Nocardiaceae** |
| 5.90 | 3.38 | 6.87 | 4.10 | 5.47 | 9.90 | 3.78 | 7.11 | 4.02 |
| ***Micromonosporaceae*** | ***Holophagaceae*** | ***Labraceae*** | ***Polyangiaceae*** | ***Rhodanobacteraceae*** | ***Weeksellaceae*** | ***Streptomycetaceae*** | ***Solirubrobacteraceae*** | |
| 3.74 | 13.3 | 8.65 | 12.4 | 11.6 | 3.80 | 3.60 | 4.37 | |

**Supplementary Table 2:** Genome-wide LOD threshold score calculated per individual taxa (phenotype) at ASV, genus and family level (1000 permutations, alpha=0.05). Source data are provided as a Source Data file.

| | | | | | | | | |
|---|---|---|---|---|---|---|---|---|
| **ASV (alpha=0.05)** | | | | | | | | |
| ***Sphingomonas env.OPS 17*** | ***Allo.Neo.Para. Rhizobium*** | ***Pedobacter1*** | ***Paenibacillus1*** | ***Niastella*** | ***Paenibacillus2*** | ***Stenotrophomonas*** | ***Paenibacillus3*** | ***Streptomyces2*** |
| 8.37 | 9.15 | 10.3 | 3.91 | 7 | 4.2 | 17.1 | 4.53 | 4.96 |
| ***Lysobacter1*** | ***Asticcacaulis*** | ***Variovorax*** | ***Paenibacillus4*** | ***Bdellovibrio*** | ***Duganella*** | ***Holophaga*** | ***Vicinamibacterales*** | ***Sphingobacteriaceae*** |
| 4.78 | 11.3 | 8.01 | 12.8 | 9.29 | 8.70 | 12.2 | 4.23 | 4.79 |
| ***Lysobacter2*** | ***Sorangium*** | ***Sphingopyxis*** | ***Chryseolinea*** | ***Tahibacter*** | ***Pedobacter2*** | ***Comamonadaceae*** | ***Chryseobacterium*** | ***Streptomyces3*** |
| 6.63 | 15.4 | 8.80 | 10.5 | 11.54 | 7.97 | 7.97 | 7.19 | 7.43 |
| ***Pedobacter3*** | ***Asticcacaulis1*** | ***Pseudomonas*** | ***Pseudomonas1*** | ***Streptomyces1*** | ***Lysobacter3*** | ***Sphingomonas*** | ***Nocardia*** | ***Streptomyces4*** |
| 12.1 | 9.80 | 7.20 | 8.63 | 6.25 | 11.6 | 7.39 | 4.71 | 11.7 |
| **Genus (alpha=0.05)** | | | | | | | | |
| ***Pedobacter*** | ***Paenibacillus*** | ***Ramlibacter*** | ***Rhodanobacter*** | ***Asticcacaulis*** | ***Thermomonas*** | ***Variovorax*** | ***Dokdonella*** | ***Chryseobacterium*** |
| 7.22 | 3.79 | 5.83 | 17.4 | 11.68 | 10.73 | 11.1 | 7.27 | 5.35 |
| ***Bdellovibrio*** | ***Microbacterium*** | ***P3OB.42*** | ***Holophaga*** | ***Sorangium*** | ***Chryseolinea*** | ***Tahibacter*** | ***Comamonadaceae*** | ***Streptomyces*** |
| 5.86 | 5.67 | 14.4 | 12.2 | 14.7 | 8.83 | 10.16 | 4.75 | 4.60 |
| **Family (alpha=0.05)** | | | | | | | | |
| ***Sphingobacteriaceae*** | ***Paenibacillaceae*** | ***Microscillaceae*** | ***Opitutaceae*** | ***Caulobacteraceae*** | ***Comamonadaceae*** | ***Chitinophagaceae*** | ***Alcaligenaceae*** | **Nocardiaceae** |
| 6.43 | 4.22 | 8.30 | 5.29 | 7.27 | 9.90 | 5.06 | 9.07 | 5.23 |
| ***Micromonosporaceae*** | ***Holophagaceae*** | ***Labraceae*** | ***Polyangiaceae*** | ***Rhodanobacteraceae*** | ***Weeksellaceae*** | ***Streptomycetaceae*** | ***Solirubrobacteraceae*** | |
| 4.54 | 14.1 | 9.20 | 13.1 | 13.4 | 5.09 | 4.60 | 6.26 | |

**Supplementary Table 3:** Confidence intervals (alpha=0.05) and percentage of variance ($R^2$) on each of the taxa showing associations with the barley genome at ASV level. Source data are provided as a Source Data file.

| ***Taxa (ASV)*** | ***Parental line*** | ***Chr.*** | ***Lower marker*** | ***Upper marker*** | ***$R^2$ within the interval per taxa*** |
|---|---|---|---|---|---|
| *Vicinamibacterales* | wild | 3H | SCRI_RS_130264 (8.8 cM) | SCRI_RS_229894 (20.4 cM) | 14.13 % |
| *Variovorax* | wild | 3H | SCRI_RS_154747 (38.75 cM) | BOPA2_12_10114 (39.4 cM) | 63.33 % |
| *Holophaga* | wild | 3H | SCRI_RS_154747 (38.75 cM) | SCRI_RS_141171 (40.6 cM) | 79.47 % |
| *Sorangium* | wild | 3H | SCRI_RS_154747 (38.75 cM) | No marker at determined genetic position (40 cM) | 25.44 % |
| *Tahibacter* | wild | 3H | SCRI_RS_154747 (38.75 cM) | SCRI_RS_141171 (40.6 cM) | 88.03 % |
| *Streptomyces4* | elite | 4H | No marker at determined genetic position (72.5 cM) | No marker at determined genetic position (77.5 cM) | 37.81 % |
| *Stenotrophomonas* | wild | 4H | SCRI_RS_188944 (94.1 cM) | No marker at determined genetic position (95 cM) | 18.46 % |
| *Vicinamibacterales* | wild | 5H | SCRI_RS_206565 (96.6 cM) | No marker determined | 13.83 % |

**Supplementary Table 4:** Confidence intervals (alpha=0.05) and percentage of variance ($R^2$) on each of the taxa showing associations with the barley genome at genus level. Source data are provided as a Source Data file.

| ***Taxa (Genus)*** | ***Parental line*** | ***Chr.*** | ***Lower marker*** | ***Upper marker*** | ***$R^2$ within the interval per taxa*** |
|---|---|---|---|---|---|
| *Ramlibacter* | wild | 2H | No marker at determined genetic position (117.5 cM) | BOPA2_12_10579 | 39.47 % |
| *Variovorax* | wild | 3H | BOPA1_2765_406 (38 cM) | SCRI_RS_141171 (40.6 cM) | 60.60 % |
| *Rhodanobacter* | wild | 3H | No marker at determined genetic position (30 cM) | SCRI_RS_154747 (38.75 cM) | 72.47 % |
| *Holophaga* | wild | 3H | SCRI_RS_154747 (38.75 cM) | SCRI_RS_141171 (40.6 cM) | 76.70 % |
| *Sorangium* | wild | 3H | SCRI_RS_154747 (38.75 cM) | No marker at determined genetic position (40 cM) | 30.65 % |
| *Tahibacter* | wild | 3H | SCRI_RS_154747 (38.75 cM) | SCRI_RS_141171 (40.6 cM) | 79.51 % |
| *Microbacterium* | wild | 5H | SCRI_RS_237352 (95.5 cM) | BOPA2_12_30867 (126.15 cM) | 35.29 % |
| *Streptomyces* | elite | 7H | BOPA1_5595_297 (133.9 cM) | SCRI_RS_6252 (140.1 cM) | 32.04 % |

**Supplementary Table 5:** Confidence intervals (alpha=0.05) and percentage of variance ($R^2$) on each of the taxa showing associations with the barley genome at family level. Source data are provided as a Source Data file.

| ***Taxa (Family)*** | ***Parental line*** | ***Chr.*** | ***Lower marker*** | ***Upper marker*** | ***$R^2$ within the interval per taxa*** |
|---|---|---|---|---|---|
| *Polyangiaceae (Sorangium)* | wild | 1H | SCRI_RS_236160 (97.9 cM) | SCRI_RS_236160 (97.9 cM) | 7.93 % |
| *Comamonaceae (Variovorax)* | wild | 3H | SCRI_RS_154747 (38.75 cM) | SCRI_RS_141171 (40.6 cM) | 67.10 % |
| *Polyangiaceae (Sorangium)* | wild | 3H | SCRI_RS_154747 (38.75 cM) | No marker at determined genetic position (40 cM) | 19.59 % |
| *Holophagaceae (Holophaga)* | wild | 3H | No marker at determined genetic position (30 cM) | SCRI_RS_141171 (40.6 cM) | 74.08 % |
| *Rhodanobacteraceae (Rhodanobacter* and *Tahibacter)* | wild | 3H | SCRI_RS_154747 (38.75 cM) | SCRI_RS_141171 (40.6 cM) | 94.26 % |
| *Sphingobacteriaceae* | wild | 5H | No marker at determined genetic position (155 cM) | SCRI_RS_167103 (161.7 cM) | 42.31 % |
| *Streptomycetaceae* | elite | 7H | BOPA1_5595_297 (133.9 cM) | SCRI_RS_6252 (140.7 cM) | 31.13 % |

**Supplementary Table 6**: Pedigree verification of the sibling lines or parental contribution in Flapjack software. Data based on at least two independent lines per genotype processed with the 50k Illumina Infinium iSelect genotyping platform.

| Genotype | Markers number | Heterozygous markers (%) | Maker identity with Barke (%) | Marker identity with a Barke x HID144 Derived (%) |
|---|---|---|---|---|
| Barke | 43,364 | 0.8 | 100 | n/a |
| 124_52 | 43,057 | 1.5 | 93.3 | 99.3 |
| 124_17 | 43,302 | 0.8 | 95.5 | 99.9 |
| HID144 | 42,833 | 0.9 | 60.3 | n/a |

**Supplementary Table 7:** Root macro architectural traits of the sibling and the elite lines. The values represent the average values (n=4) the standard error of the mean (SEM). Differences were tested using a Kruskal test or ANOVA, p-value<0.05. Source data are provided as a Source Data file.

| *Genotype* | *Dry shoot weight (g)* | *Dry root weight (g)* | *Primary root length (cm)* | *Surface area ($cm^2$)* | *Average diameter (mm)* | *Root Volume ($cm^3$)* | *Specific root length* | *Root density ($g/cm^3$)* | *Tips* | *Forks* | *Crosslinks* |
|---|---|---|---|---|---|---|---|---|---|---|---|
| **Elite** | 0.32 ±0.05 | 0.29 ±0.17 | 48 ±5 | 291 ±7 | 2.2 ±0.4 | 18 ±6 | 2804 ±0.01 | 0.022 ±0.009 | 1475 ±3 | 4328 ±9 | 129 ±71 |
| **124_17** | 0.38 ±0.05 | 0.34 ±0.19 | 44 ±6 | 387 ±35 | 3.0 ±0.4 | 30 ±6 | 2212 ±676 | 0.018 ±0.013 | 1481 ±2 | 4874 ±6 | 113 ±26 |
| **124_52** | 0.37 ±0.03 | 0.13 ±0.01 | 41 ±2 | 312 ±31 | 2.1 ±0.3 | 17 ±4 | 3831 ±348 | 0.009 ±0.002 | 1204 ±2 | 4011 ±8 | 105 ±27 |

**Supplementary Table 8**: RNA-seq metadata table. Brep stand for biological replicate, while srep is sequencing replicate.

| GENOTYPE | BREP | SREP | SAMPLING DATE | *QRMC-3HS* LOCUS GENO TYPE | QUANT FILES |
|---|---|---|---|---|---|
| **124_17** | brep1 | srep1 | 08/05/2019 | Elite | CEM01_JH15-105 |
| **124_17** | brep2 | srep1 | 06/05/2019 | Elite | CEM01_JH15-14 |
| **124_17** | brep3 | srep1 | 06/05/2019 | Elite | CEM01_JH15-26 |
| **124_17** | brep4 | srep1 | 06/05/2019 | Elite | CEM01_JH15-51 |
| **124_17** | brep5 | srep1 | 06/05/2019 | Elite | CEM01_JH15-7 |
| **124_52** | brep1 | srep1 | 08/05/2019 | Wild | CEM01_JH15-104 |
| **124_52** | brep2 | srep1 | 08/05/2019 | Wild | CEM01_JH15-115 |
| **124_52** | brep3 | srep1 | 06/05/2019 | Wild | CEM01_JH15-32 |
| **124_52** | brep4 | srep1 | 06/05/2019 | Wild | CEM01_JH15-35 |
| **124_52** | brep5 | srep1 | 07/05/2019 | Wild | CEM01_JH15-95 |
| **BARKE** | brep1 | srep1 | 08/05/2019 | Elite | CEM01_JH15-113 |
| **BARKE** | brep2 | srep1 | 08/05/2019 | Elite | CEM01_JH15-117 |
| **BARKE** | brep3 | srep1 | 06/05/2019 | Elite | CEM01_JH15-22 |
| **BARKE** | brep4 | srep1 | 07/05/2019 | Elite | CEM01_JH15-91 |
| **BARKE** | brep5 | srep1 | 07/05/2019 | Elite | CEM01_JH15-96 |

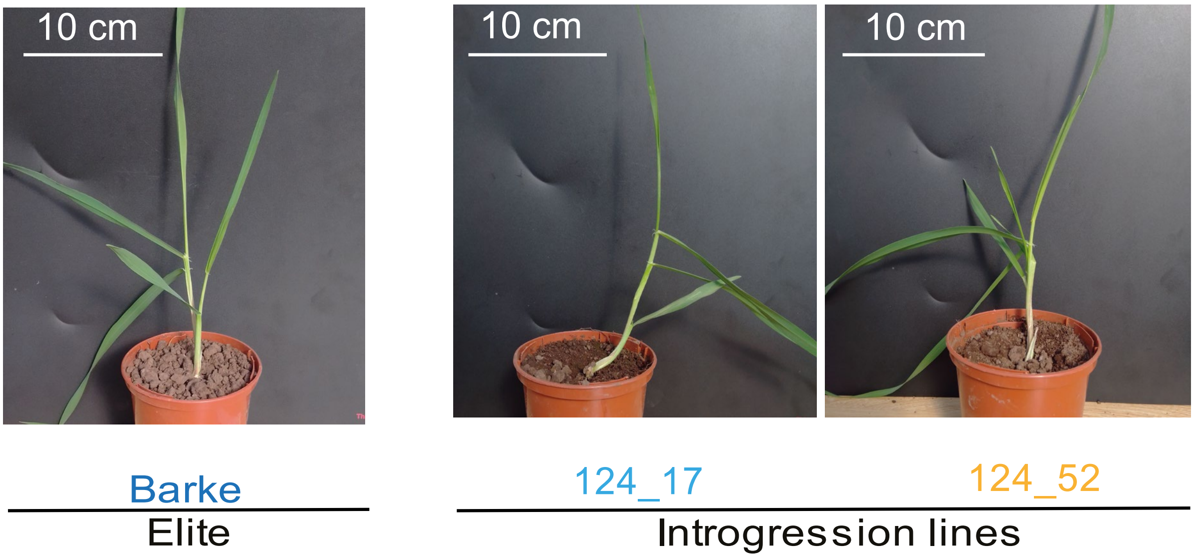


**Supplementary Fig. 1:** Pictures of the sibling lines 124_52, 124_17 and the elite genotype Barke at elongation stem stage when their rhizosphere microbiota is harvested.

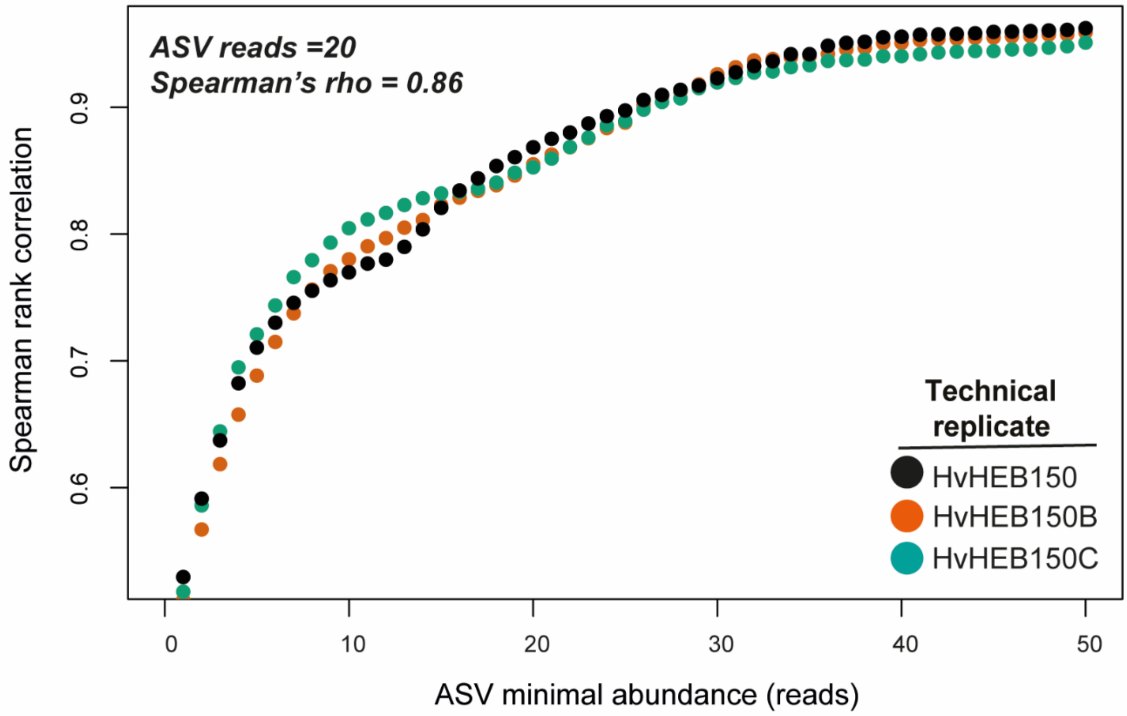


**Supplementary Fig. 2:** Technical replicate pairwise correlations to assess technical reproducibility at minimum sample size for downstream reads filtering. Source data are provided as a Source Data file.

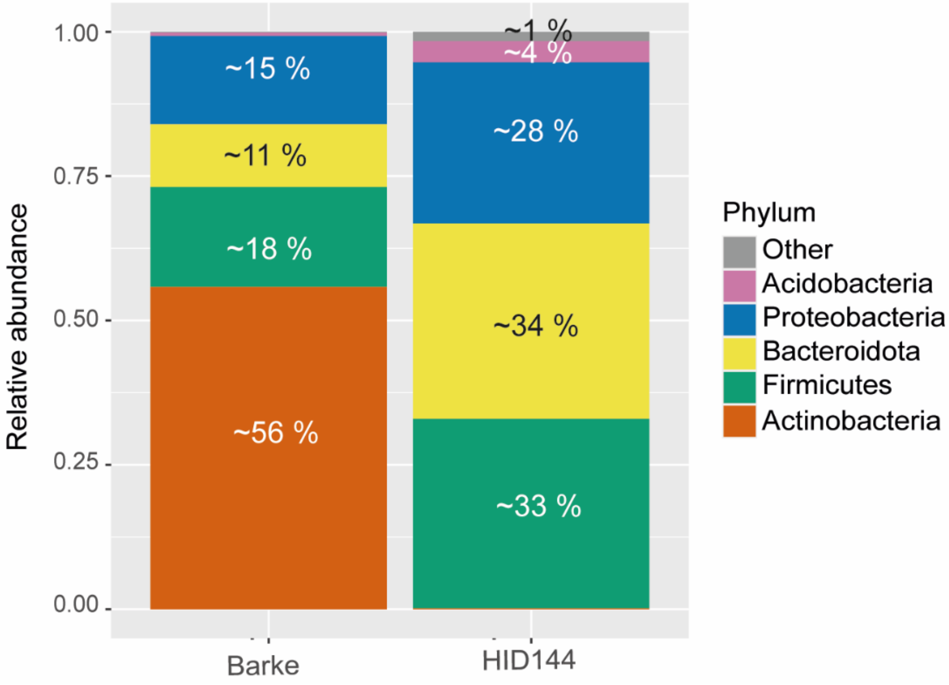


**Supplementary Fig. 3:** Stacked bar plots showing the main phyla differentially abundant between the parental lines Barke (elite) and HID144 (wild). Source data are provided as a Source Data file.

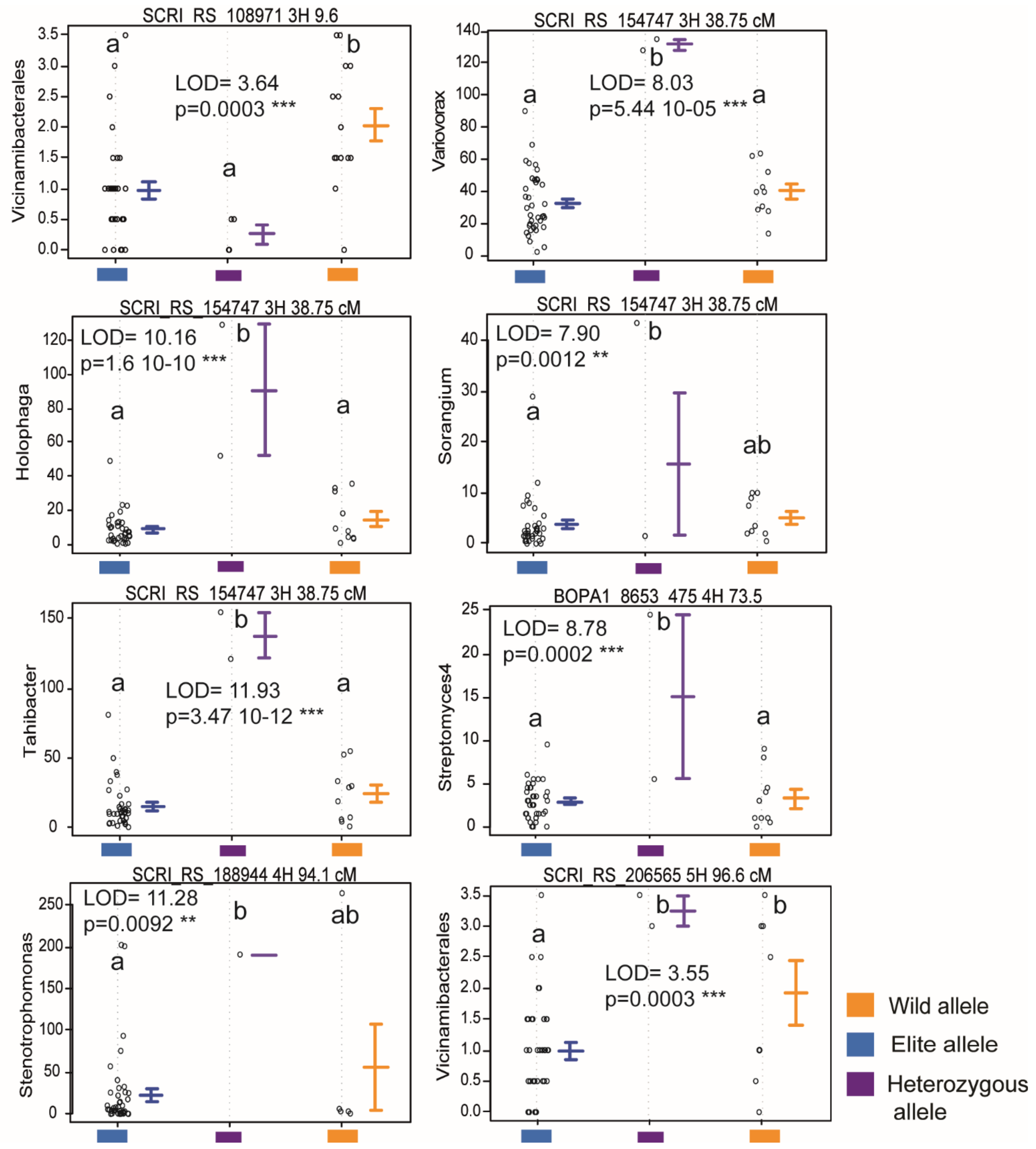


**Supplementary Fig. 4:** Bacterial abundances in sequencing reads (ASV level, y-axis) in members of the population. Individual dots depict individual biological replicates color-coded according to allelic composition at molecular markers indicated at the top of each panel with chromosomal location (e.g., 3H, chromosome 3H) and map position in centimorgan (cM). Coloured bars indicate the phenotypic mean values with +/- the standard error. Different lowercase letters denote significant differences calculated using ANOVA post-hoc Tukey or Kruskal–Wallis and post-hoc Dunn's test, at the indicated *P*-values. LOD, $Log_{10}$ likelihood ratio of a QTL presence at that marker position. Source data are provided as a Source Data file.

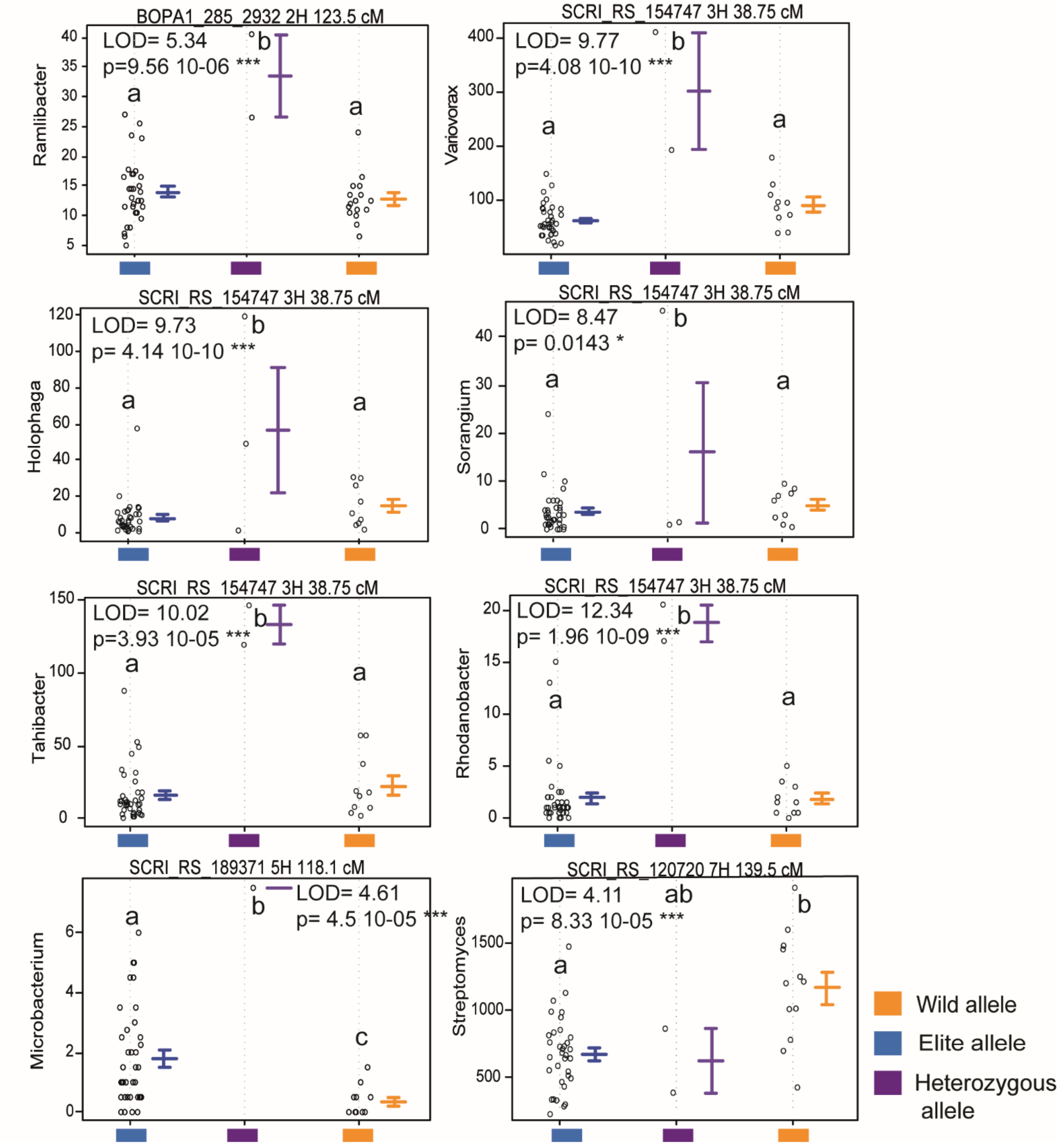


**Supplementary Fig. 5:** Bacterial abundances in sequencing reads (genus level, y-axis) in members of the population. Individual dots depict individual biological replicates color-coded according to allelic composition at molecular markers indicated at the top of each panel with chromosomal location (e.g., 3H, chromosome 3H) and map position in centimorgan (cM). Coloured bars indicate the phenotypic mean values with +/- the standard error. Different lowercase letters denote significant differences calculated using ANOVA post-hoc Tukey or Kruskal–Wallis and post-hoc Dunn's test, at the indicated *P*-values. LOD, $Log_{10}$ likelihood ratio of a QTL presence at that marker position. Source data are provided as a Source Data file.

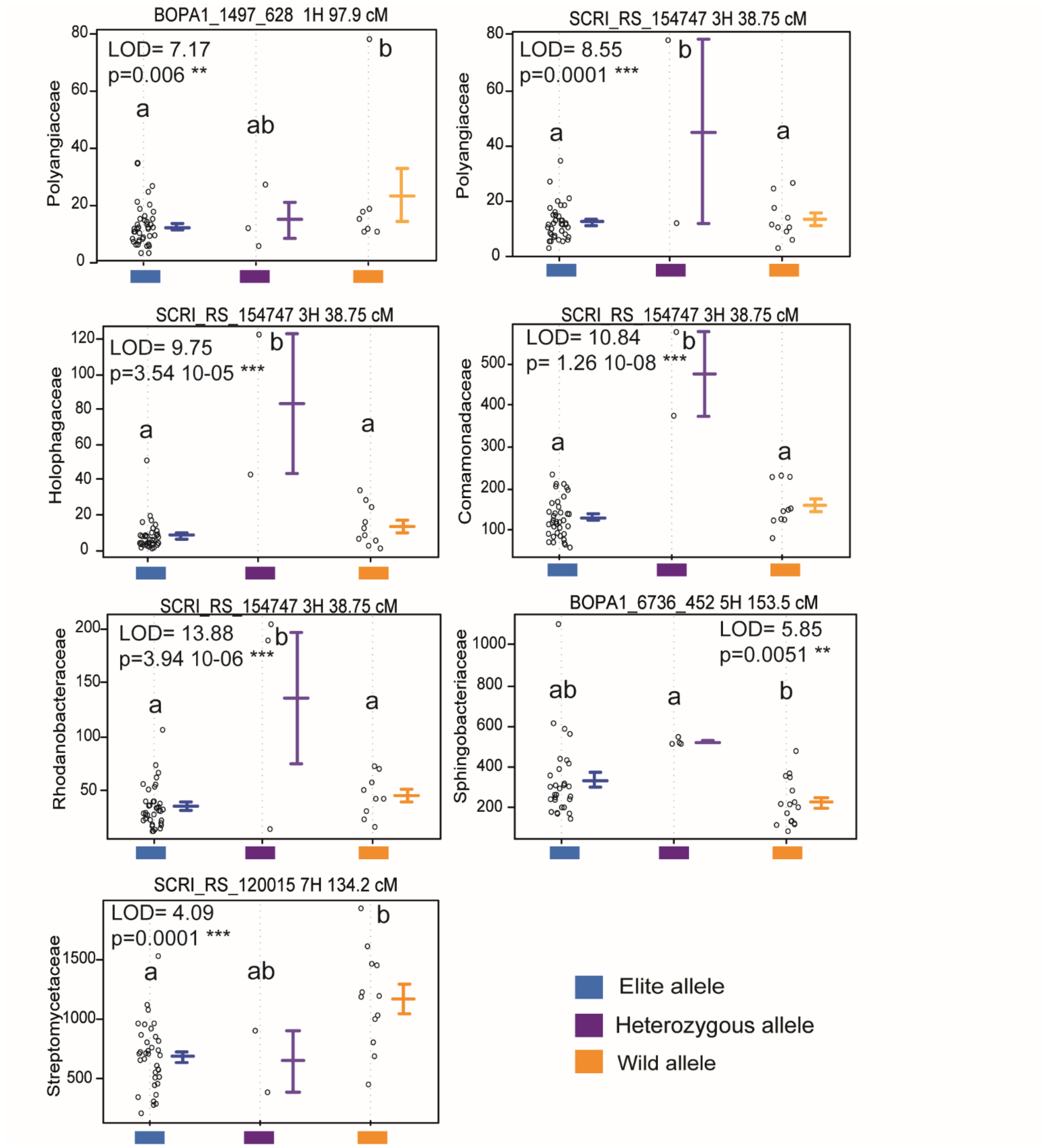


**Supplementary Fig. 6:** Bacterial abundances in sequencing reads (familylevel, y-axis) in members of the population. Individual dots depict individual biological replicates color-coded according to allelic composition at molecular markers indicated at the top of each panel with chromosomal location (e.g., 3H, chromosome 3H) and map position in centimorgan (cM). Coloured bars indicate the phenotypic mean values with +/- the standard error. Different lowercase letters denote significant differences calculated using ANOVA post-hoc Tukey or Kruskal–Wallis and post-hoc Dunn's test, at the indicated *P*-values. LOD, $Log_{10}$ likelihood ratio of a QTL presence at that marker position. Source data are provided as a Source Data file.

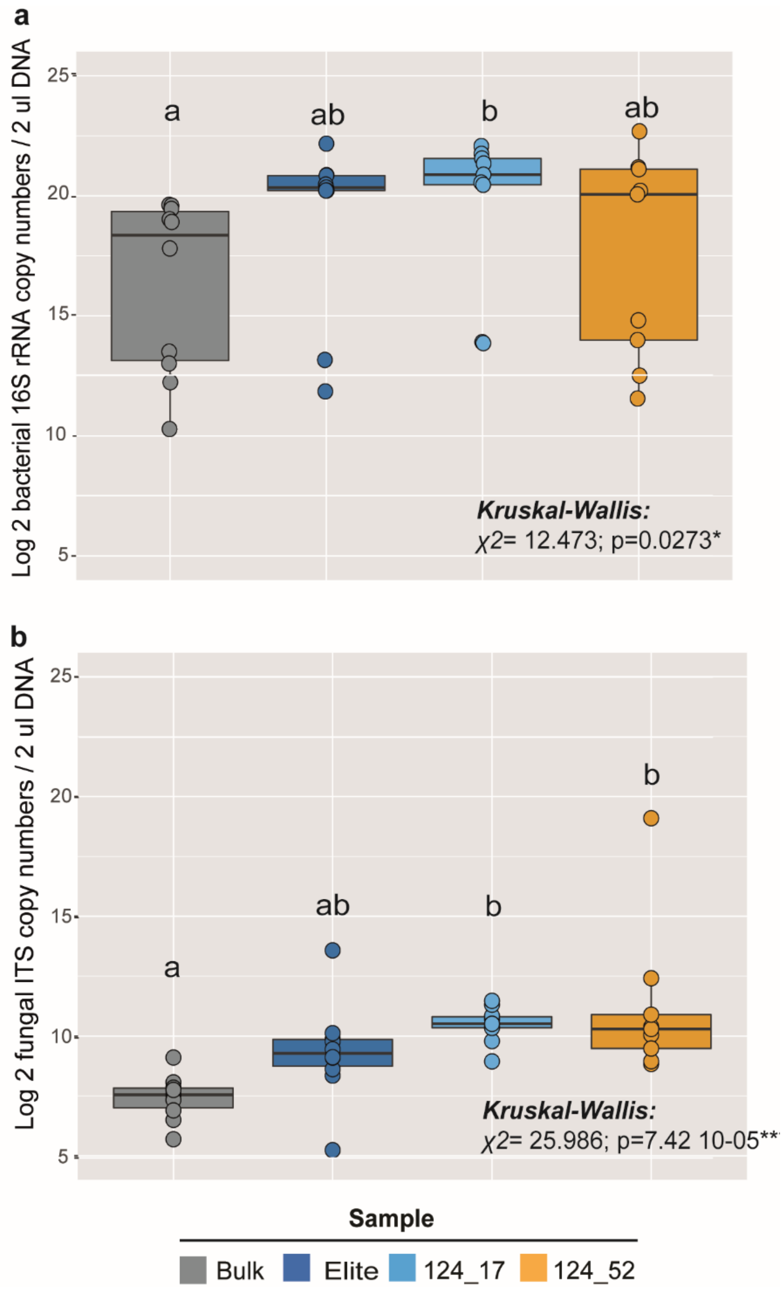


**Supplementary Fig. 7:** Quantification of bacterial and fungal DNA in the sibling lines, the elite genotype and bulk soil. **a)** Boxplots depicting the logarithm (base 2) of the number of 16S copies per ul of sample DNA. **b)** Boxplots illustrating the logarithm (base 2) of the number of ITS copies per ul of sample DNA. In each panel, individual dots depict individual biological replicates. Upper and lower edges of the box plots represent the upper and lower quartiles, respectively. The bold line within the box denotes the median. Whiskers denote values within 1.5 interquartile ranges. Different letters indicate significantly different samples (Kruskal–Wallis and post-hoc Dunn's test, at the indicate P values). Source data are provided as a Source Data file.

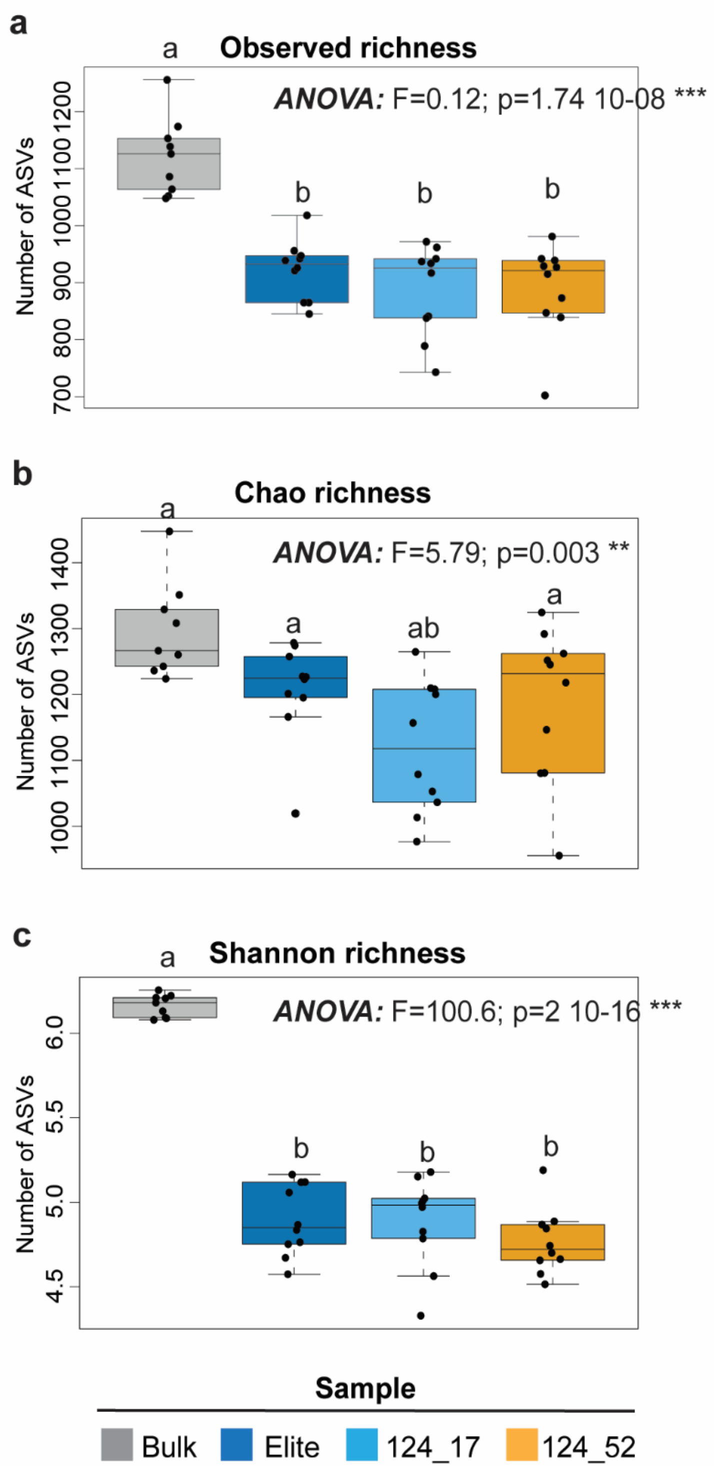


**Supplementary Fig. 8:** Boxplots depicting alpha-diversity indexes. **a)** and **b)** show the community richness (observed species, Chao1) and c) diversity (Shannon) for the unplanted soil, elite and the sibling lines genotypes. In each panel, individual dots depict individual biological replicates. Upper and lower edges of the box plots represent the upper and lower quartiles, respectively. The bold line within the box denotes the median. Whiskers denote values within 1.5 interquartile ranges. Letters indicate significant differences following ANOVA and post-hoc Tukey test as indicated in the individual panels. Source data are provided as a Source Data file

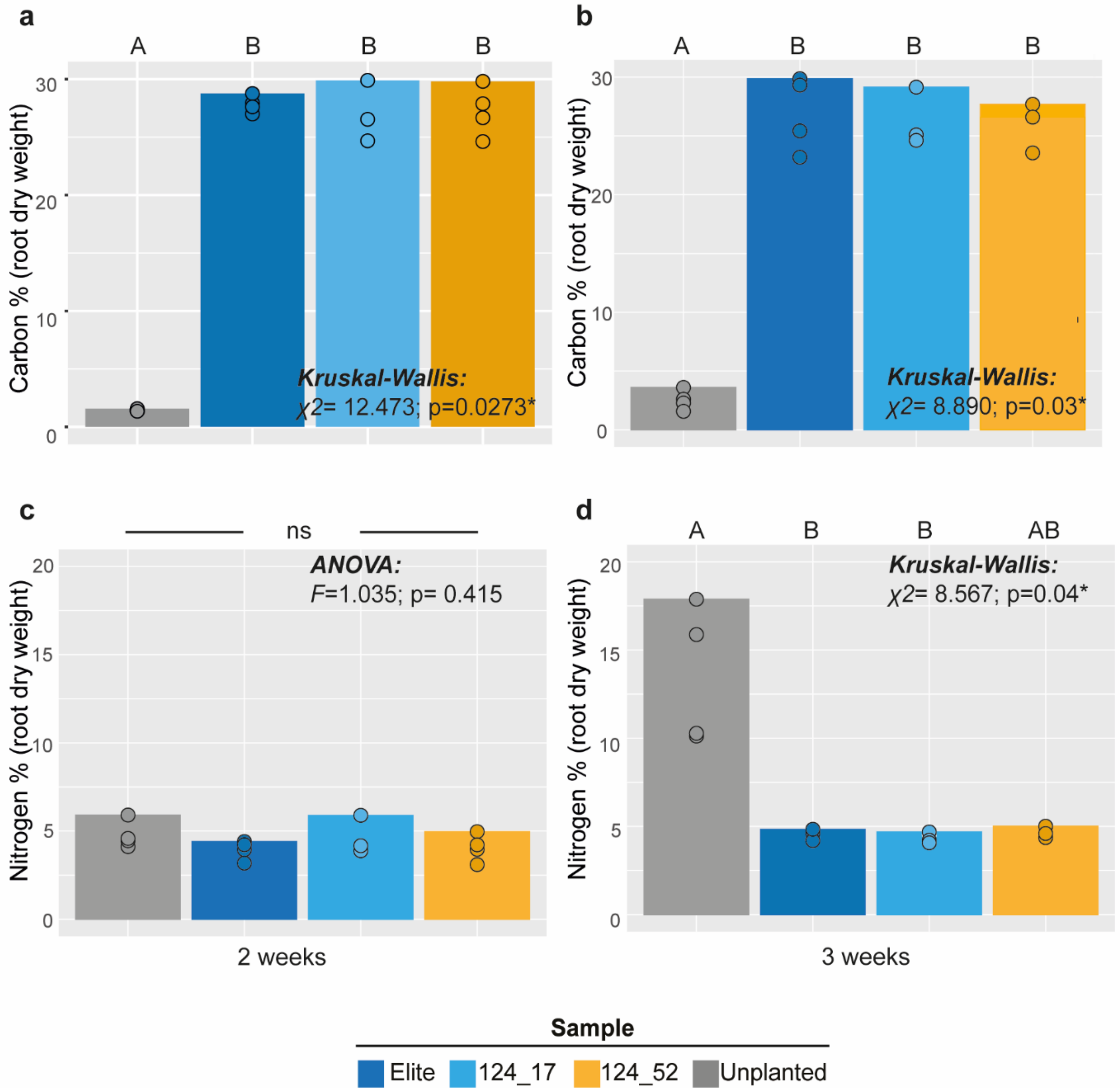


**Supplementary Fig. 9:** Bar plots representing the mean carbon and nitrogen content (% per weight) in the exudates of the sibling lines, the elite genotypes exudates and the unplanted control at two timepoints. Individual dots depict biological replicates. **a)** Carbon content at 2 weeks timepoint. **b)** Carbon content at 3 weeks timepoint. **c)** Nitrogen content at 2 weeks timepoint. **d)** Nitrogen content at 3 weeks timepoint. In each panel, the upper edge of the box depicts mean value, individual dots are individual biological replicates. Uppercase letters denote significant differences at the indicated statistic; ns no significant differences at the imposed threshold. Source data are provided as a Source Data file.

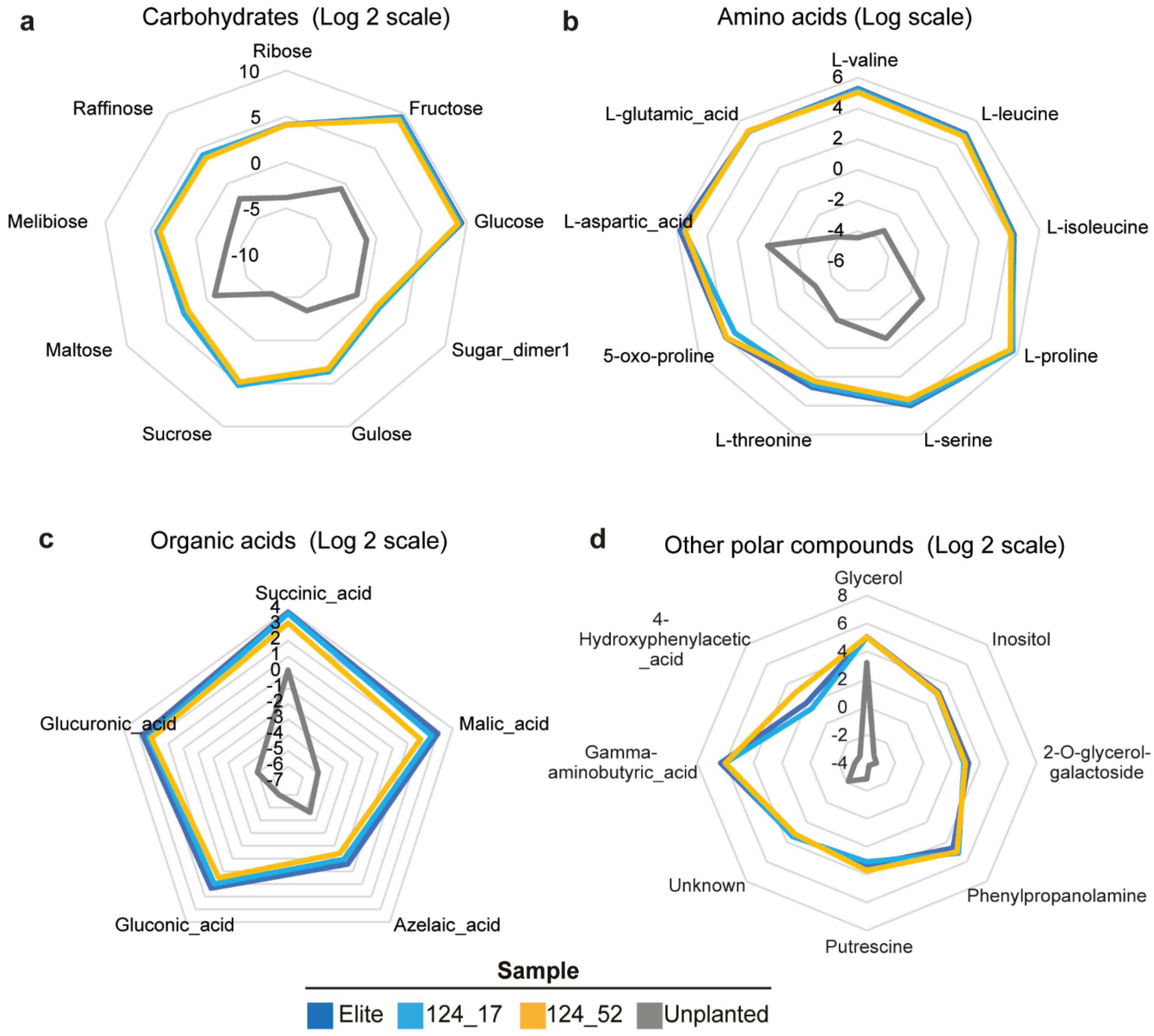


**Supplementary Fig. 10:** Radar plots depicting the primary metabolism of root exudates and unplanted controls. Different panels indicate categories of compounds retrieved from root exudates **a)** carbohydrates, **b)** amino acids, **c)** acids and **d)** other polar compounds. Yellow, blue and light blue lines represent different plant genotypes exudates composition. Grey lines indicate the unplanted control. Individual numbers depict $log_2$ scaled to the means of four blocks representing cumulative 60 biological replicates per genotype. Source data are provided as a Source Data file.

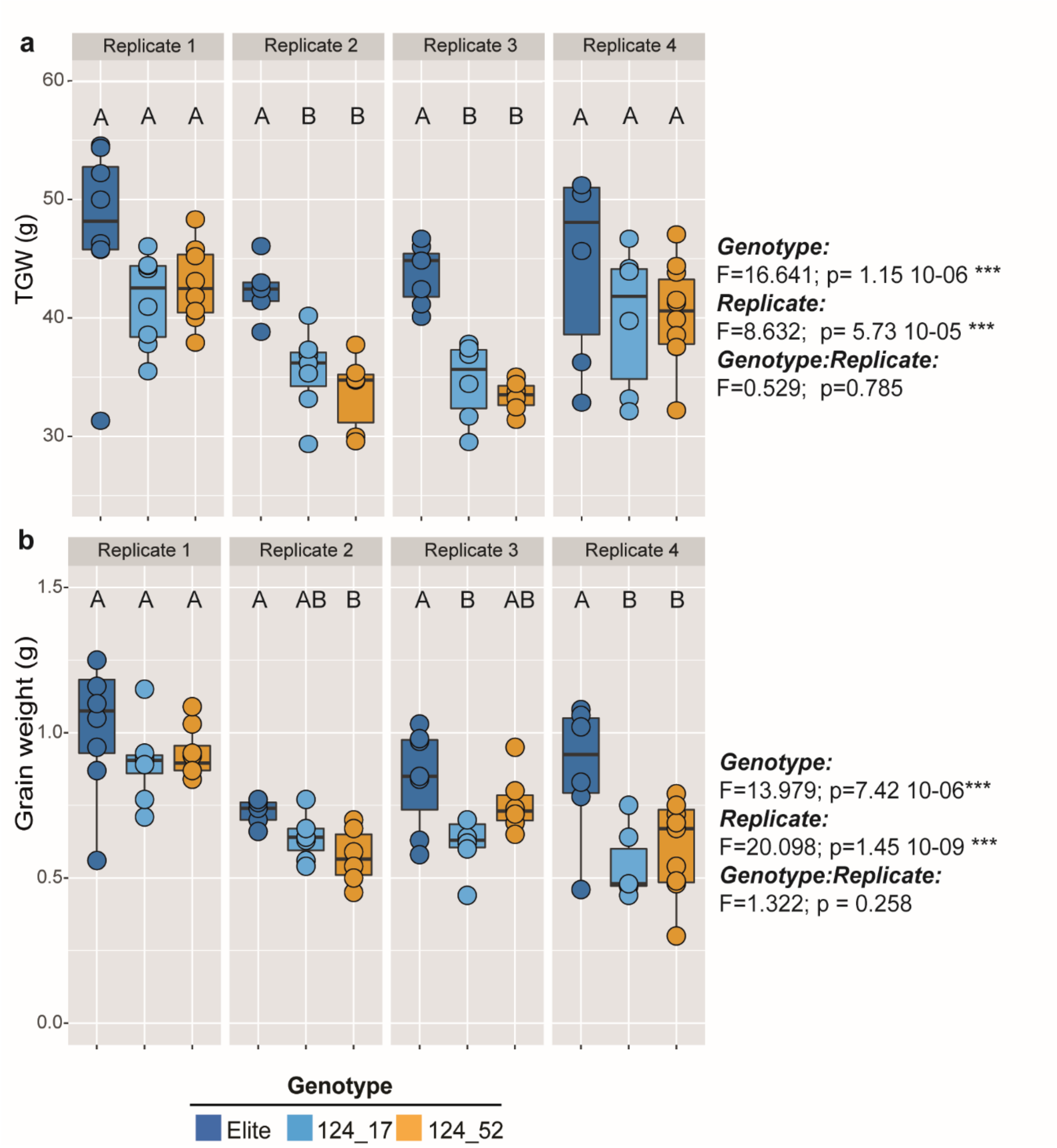


**Supplementary Fig. 11:** Boxplot representing different barley yield parameters. **a)** thousand grain weight and **b)** total yield (in grams) per main tiller in the sibling lines and the elite genotype Barke. In each panel, data is shown in four independent experiments to observe the significant effect of the replication. Individual dots depict individual biological replicates. Upper and lower edges of the box plots represent the upper and lower quartiles, respectively. The bold line within the box denotes the median. Whiskers denote values within 1.5 interquartile ranges. Uppercase letters denote significant differences determined using an ANOVA followed by post-hoc Tukey test for the factors as indicated at the side of the panel. Source data are provided as a Source Data file.

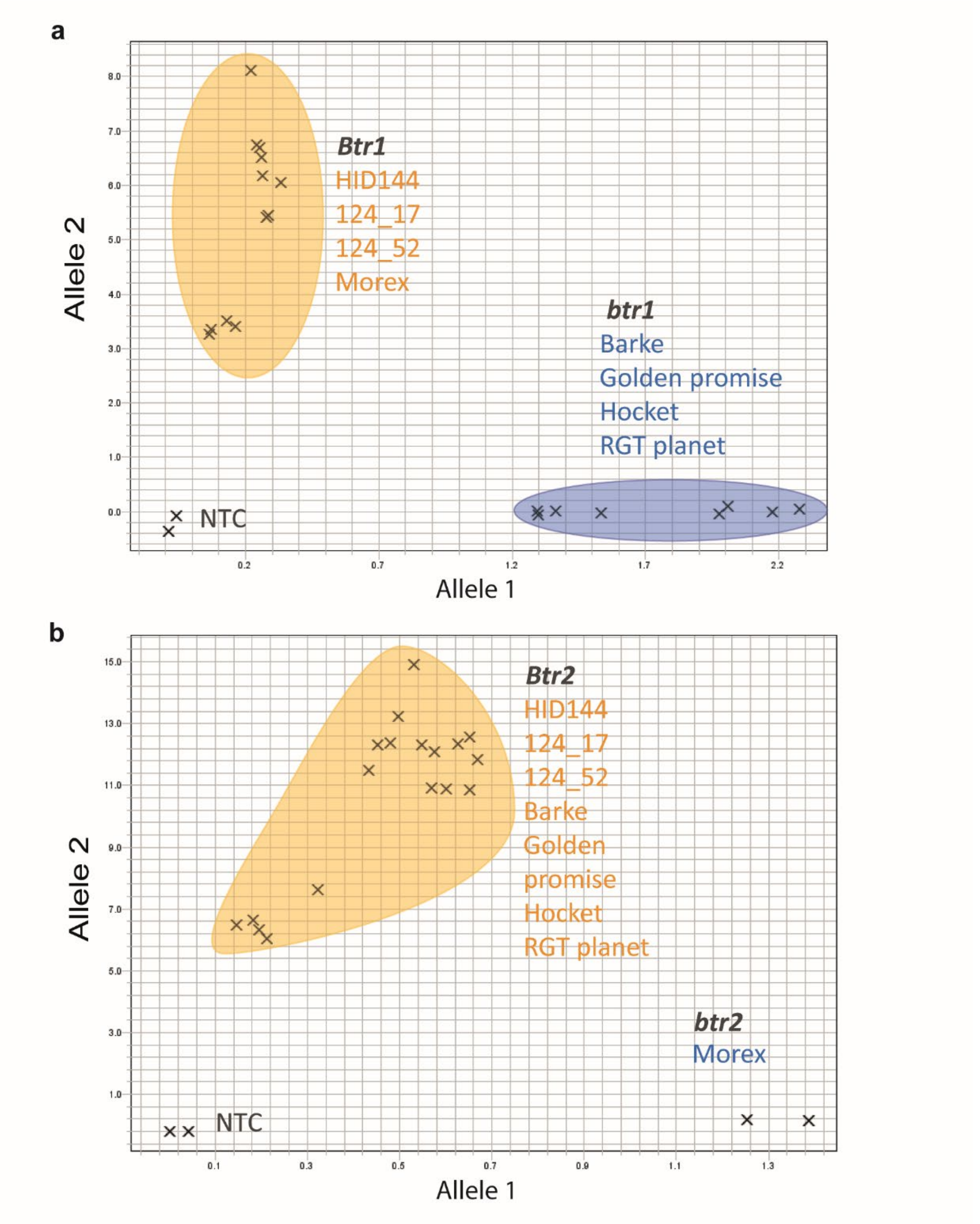


**Supplementary Fig. 12:** Allele discrimination scatter plot for mutations in the brittle rachis genes **a)** Btr1 and **b)** Btr2 in the genotypes HID144, Barke, 124_17, 124_52, Morex, Golden Promise, Hockett and RGT planet. The x-axis in **a)** represents the relative fluorescent emission for the C allele-specific probe and the y-axis represents the represents the relative fluorescent emission for the G Allele. The x-axis in **b)** represents the relative fluorescent emission for the T allele-specific probe and the y-axis represents the represents the relative fluorescent emission for the G Allele. NTC represents no template control. Source data are provided as a Source Data file.

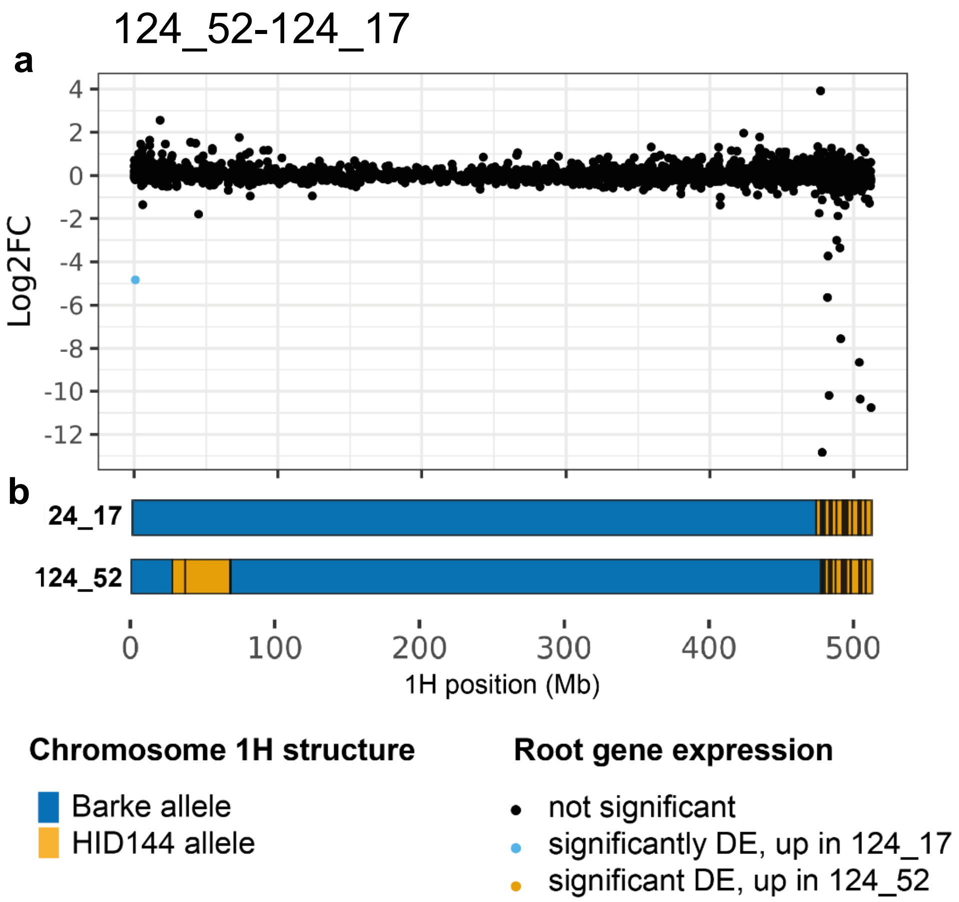


**Supplementary Fig. 13:** **a)** $log_2$ fold changes of genes with their position on barley chromosome 1H, compared to **b)** allelic information for lines 124_17 and 124_52. HID-144 introgressions are illustrated in yellow, Barke background is shown in blue. Genes are coloured according to significance, with non-DE genes coloured in black, those genes upregulated in 124_52 when compared to 124_17 in yellow, and those downregulated in the same comparison in blue. Source data are provided as a Source Data file.

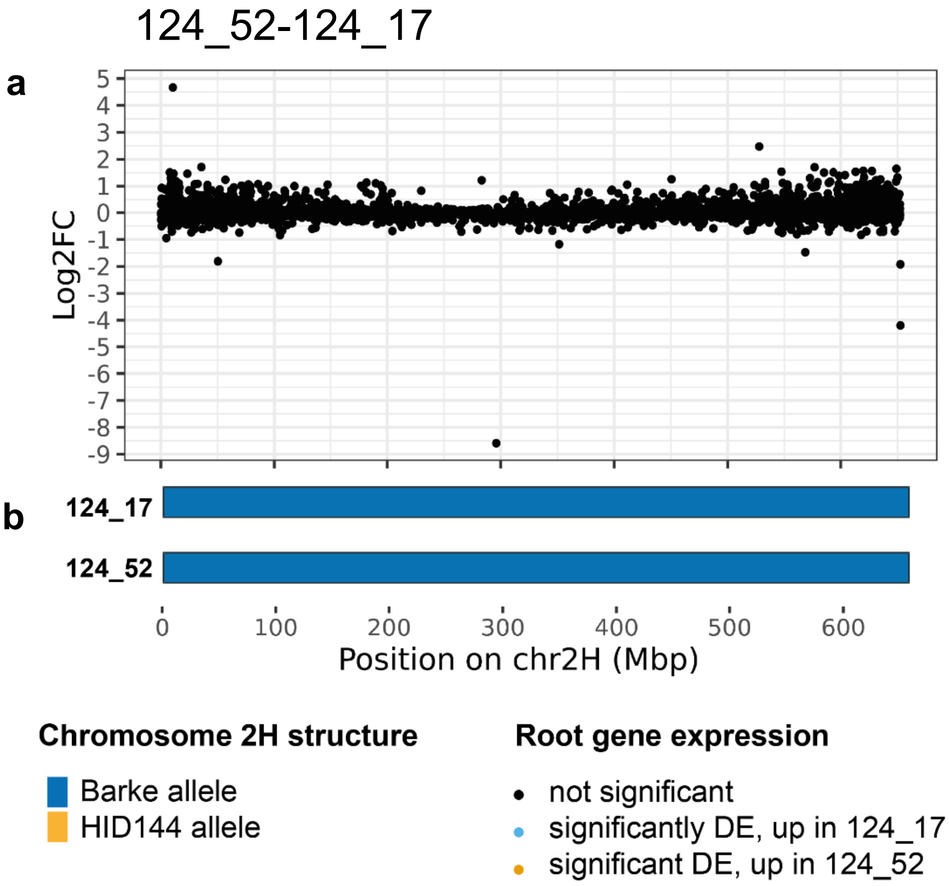


**Supplementary Fig. 14:** **a)** $log_2$ fold changes of genes with their position on barley chromosome 2H, compared to **b)** allelic information for lines 124_17 and 124_52. HID-144 introgressions are illustrated in yellow, Barke background is shown in blue. Genes are coloured according to significance, with non-DE genes coloured in black, those genes upregulated in 124_52 when compared to 124_17 in yellow, and those downregulated in the same comparison in blue. Source data are provided as a Source Data file.

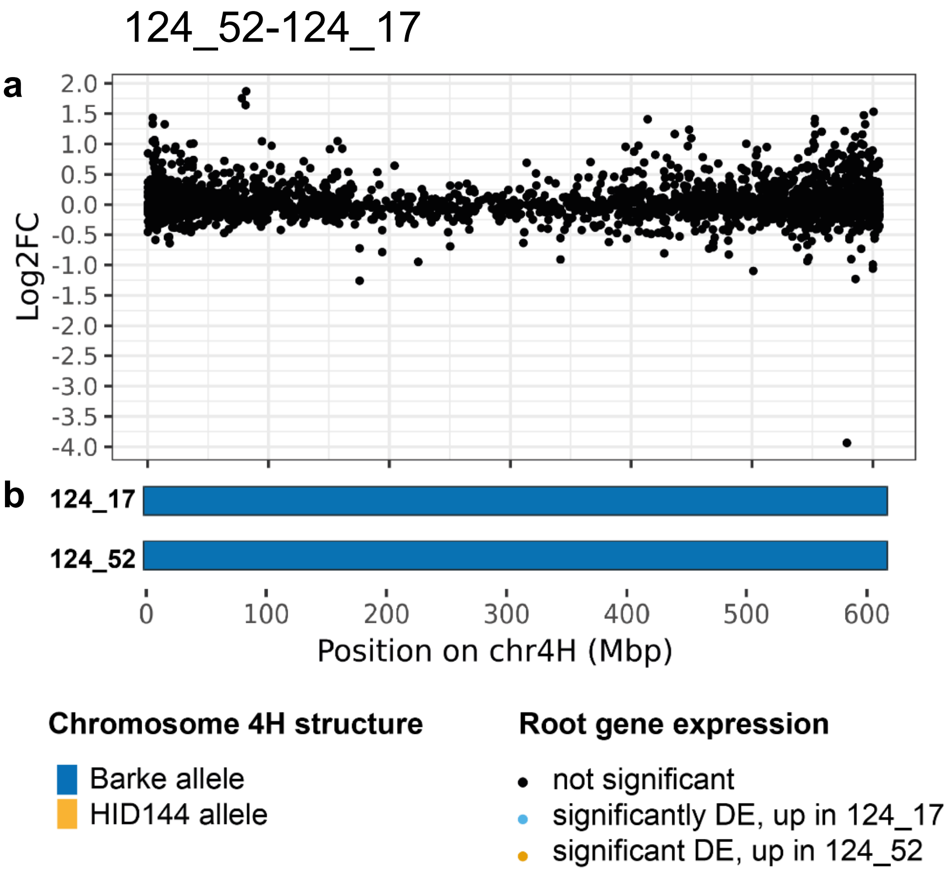


**Supplementary Fig. 15:** **a)** $\log_2$ fold changes of genes with their position on barley chromosome 4H, compared to **b)** allelic information for lines 124_17 and 124_52. HID-144 introgressions are illustrated in yellow, Barke background is shown in blue. Genes are coloured according to significance, with non-DE genes coloured in black, those genes upregulated in 124_52 when compared to 124_17 in yellow, and those downregulated in the same comparison in blue. Source data are provided as a Source Data file.

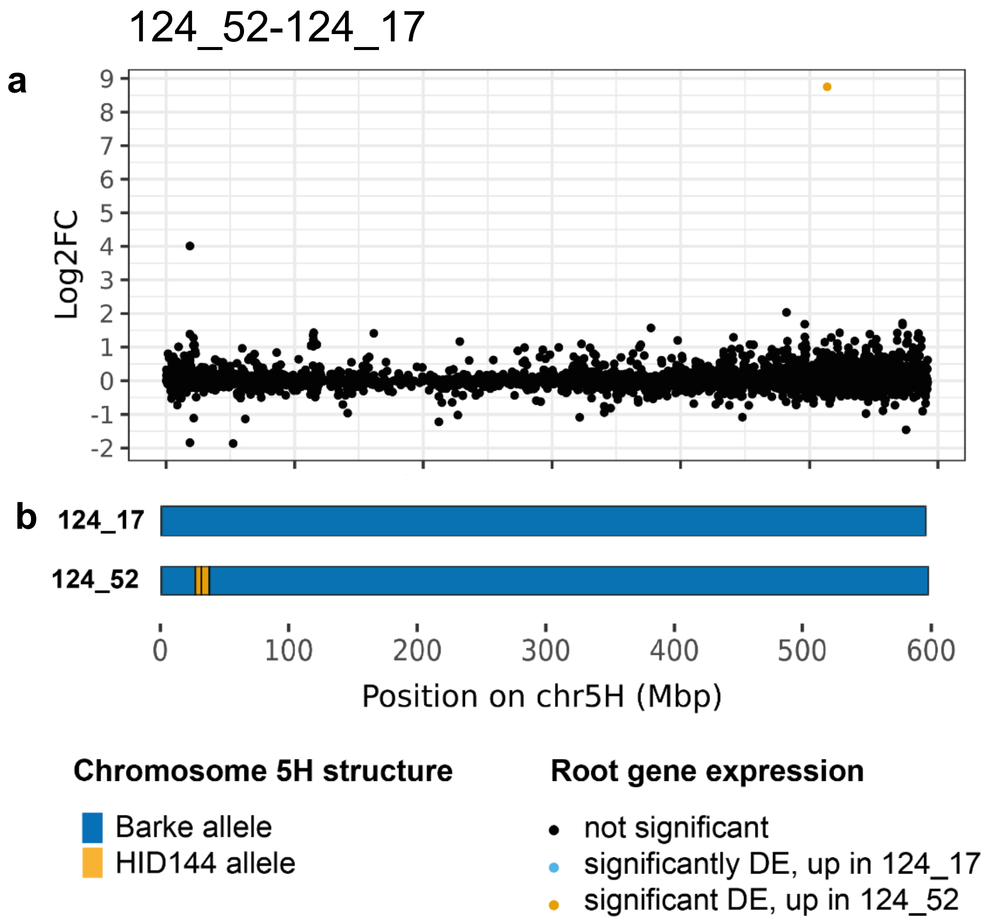


**Supplementary Fig. 16:** **a)** $\log_2$ fold changes of genes with their position on barley chromosome 5H, compared to **b)** allelic information for lines 124_17 and 124_52. HID-144 introgressions are illustrated in yellow, Barke background is shown in blue. Genes are coloured according to significance, with non-DE genes coloured in black, those genes upregulated in 124_52 when compared to 124_17 in yellow, and those downregulated in the same comparison in blue. Source data are provided as a Source Data file.

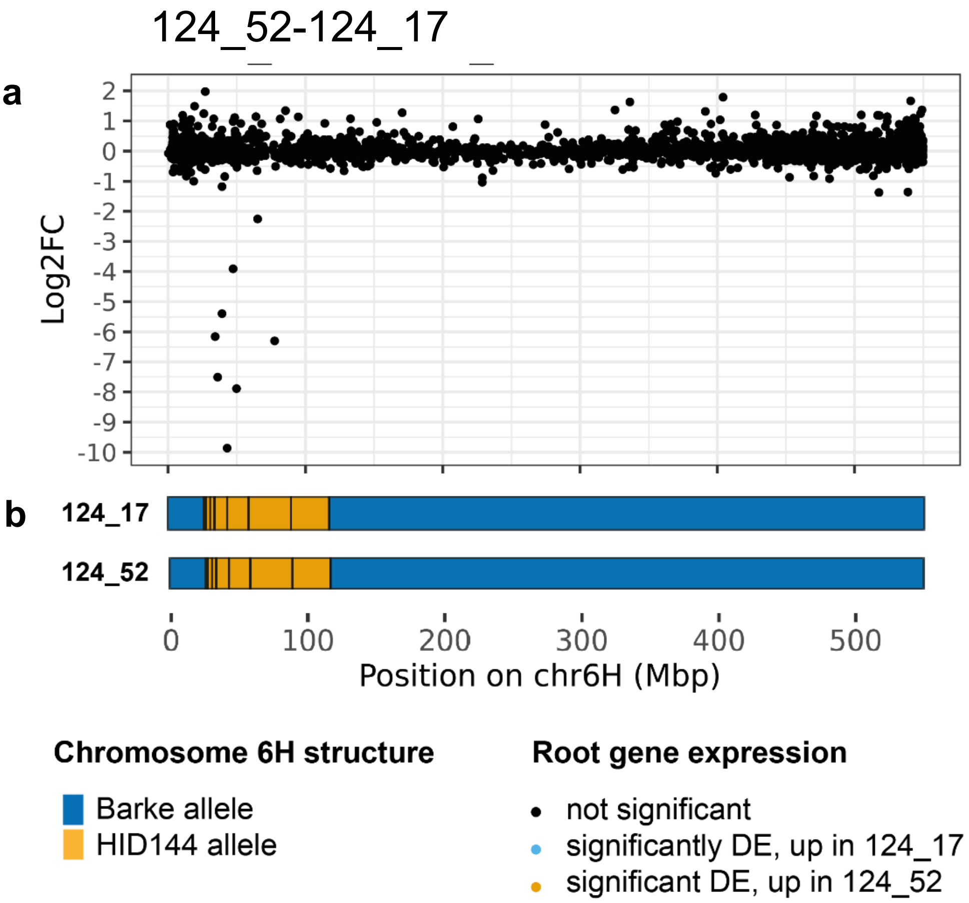


**Supplementary Fig. 17:** **a)** $\log_2$ fold changes of genes with their position on barley chromosome 6H, compared to **b)** allelic information for lines 124_17 and 124_52. HID-144 introgressions are illustrated in yellow, Barke background is shown in blue. Genes are coloured according to significance, with non-DE genes coloured in black, those genes upregulated in 124_52 when compared to 124_17 in yellow, and those downregulated in the same comparison in blue. Source data are provided as a Source Data file.

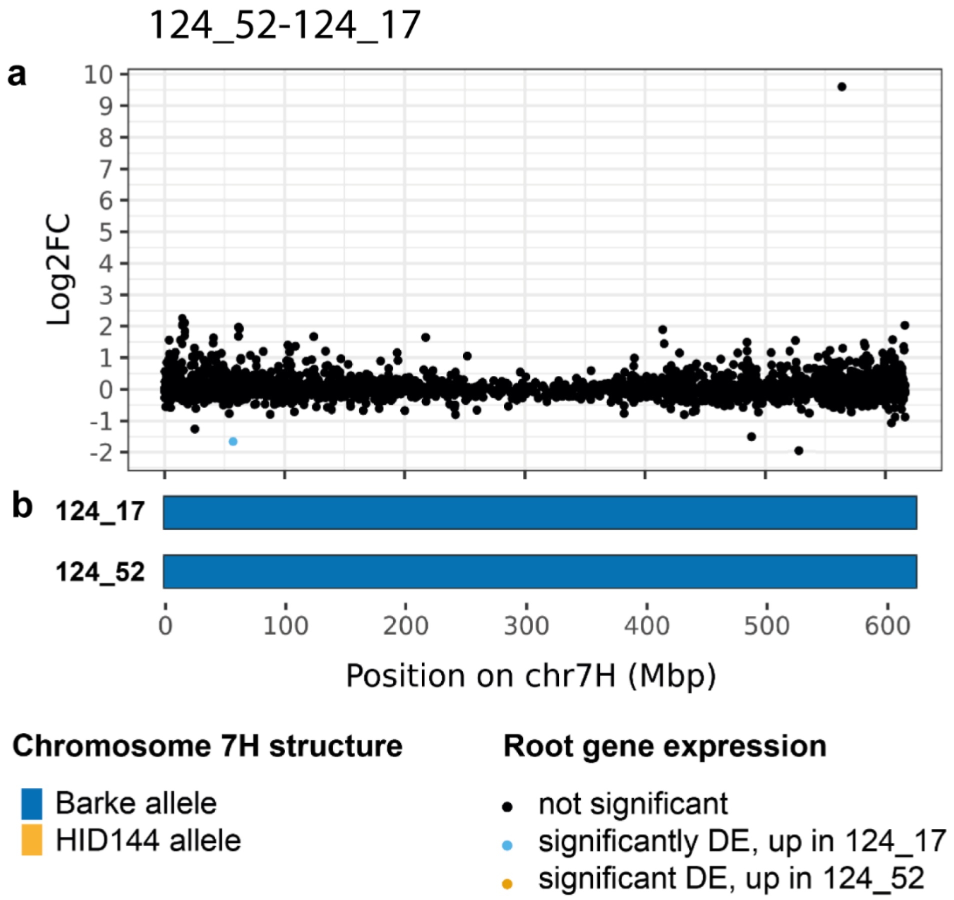


**Supplementary Fig. 18:** **a)** $\log_2$ fold changes of genes with their position on barley chromosome 7H, compared to **b)** allelic information for lines 124_17 and 124_52. HID-144 introgressions are illustrated in yellow, Barke background is shown in blue. Genes are coloured according to significance, with non-DE genes coloured in black, those genes upregulated in 124_52 when compared to 124_17 in yellow, and those downregulated in the same comparison in blue. Source data are provided as a Source Data file.

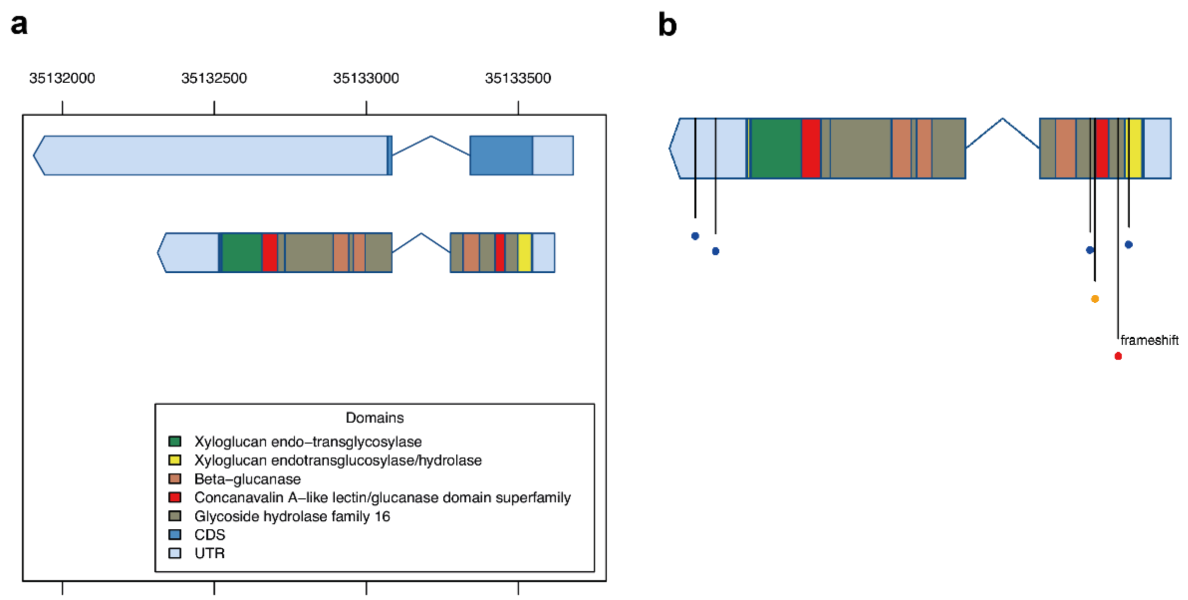


**Supplementary Fig. 19**: Schematics of transcripts in gene BaRT2v18chr3HG123140 annotated as a xyloglucan endotransglucosylase/hydrolase, illustrating results of variant calling and annotation. **a)** Two transcripts present in BaRT2v18chr3HG123140, annotated with domains using InterProScan. **b)** The second productive transcript annotated with variants. Low impact variants (those not affecting predicted amino acid sequence) are shown in blue, moderate impact variants (those causing localised amino acid changes) are shown in yellow, and a high impact frameshift variant is shown in red. Source data are provided as a Source Data file.

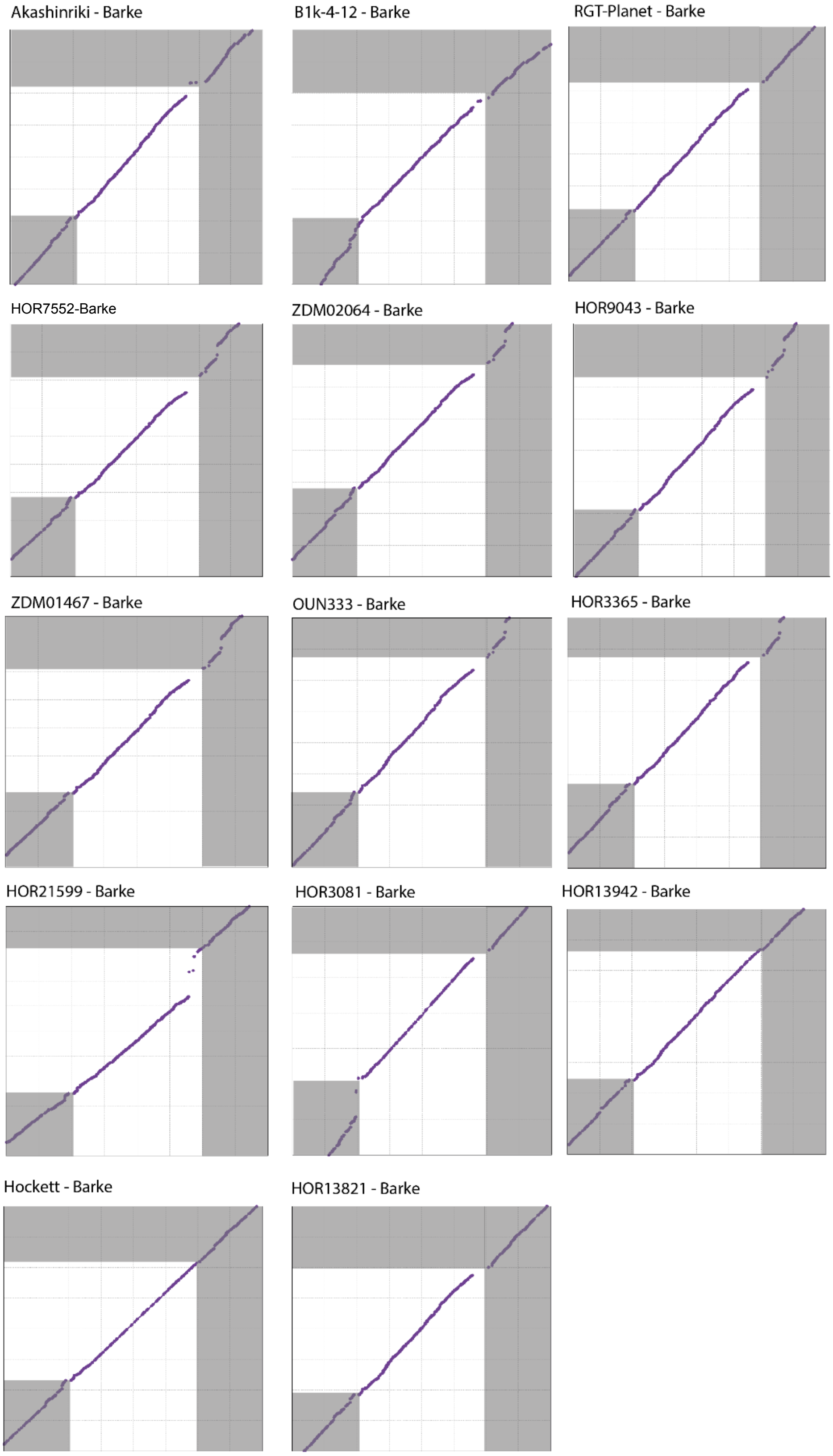


**Supplementary Fig. 20**: MUMmer plots showing collinearity of sequence between Barke (x-axis) and 14 other pan-genome genotypes (y-axis) in and around the *QRMC-3HS* locus. Area in white shows the position of the *QRMC-3HS* locus, grey areas are outside the *QRMC-3HS*. Purple dots are sequence matches between two genotypes. Source data available at Source data are provided as a Source Data file.

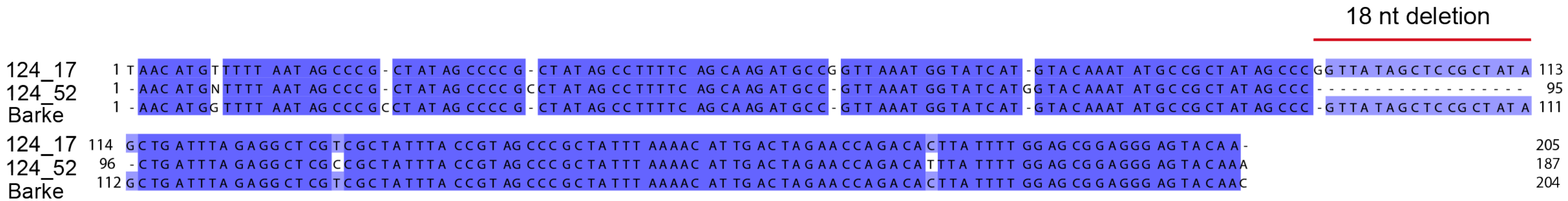


**Supplementary Fig. 21**: Nucleotide alignment showing the NLR diagnostic marker amplicon sequences (18 nt deletion flanking regions). Sanger sequences of the NLR marker amplicon were used as input. The alignment was generated in Jalview 2.11.2.2 using the Muscle alignment with default parameters. The level of sequence similarity is indicated by dark (high identity) to light purple colour (low identity). Source data are provided as a Source Data file.
